# Evidence for a vocal signature in the rat and its reinforcing effects

**DOI:** 10.1101/2021.06.07.447373

**Authors:** Cassandre Vielle, Christian Montanari, Yann Pelloux, Christelle Baunez

## Abstract

While the term “language” is used for human and non-human primates, “vocal communication” is rather used for rodents or other species. The main difference is that there is, to date, no evidence for a vocal signature in the well-known 50- and 22-kHz ultrasonic vocalizations (USV) emitted by rats. Here, we show that rats can recognize the identity of the USV emitter since they self-administer preferentially playback of 50-kHz USV emitted by a stranger rat over those emitted by their cage-mate. In a second experiment, we show that the familiarity with the USV emitter also modulate the effect of USV playback during cocaine self-administration, since only stranger, but not familiar, 50-kHz USV decrease drug intake. Finally, to study the neurobiological substrate of those processes, we have tested the effects of the subthalamic nucleus (STN) lesion on these various conditions. STN-lesioned rats did not lever press much for any USV playback, whatever their emotional valence, nor did they seem able to differentiate familiar from stranger peer. Advocating for the existence of a vocal signature in rats, these results highlight the importance of ultrasonic communication in socio-affective influence of behavior, such as the influence of proximal social factors on drug consumption and confirm the role of the subthalamic nucleus on this influence.

## Introduction

Like humans, rats are highly social animals, living in complex social groups (for review: Barnett, 1967; Lund, 1975) and display a large panel of socio-affective skills (from emotion recognition, to emotion contagion, and empathy; for review Schweinfurth, 2020). These abilities make rodent models relevant to study the influence of socio-affective factors on behavior. For instance, studies on drug consumption showed that the presence of a peer modulates the drug intake (Smith, 2012), in a substance-specific way (Gipson et al., 2011). In both rats and humans, the presence of a peer decreases cocaine intake (Giorla et al., 2018), while when given the choice, young rats favor social access over methamphetamine in young rats (Venniro et al., 2018). The protective role of positive social contacts over drug consumption is mostly interpreted as an alternative, that overcomes the rewarding value of drugs (Fritz et al., 2011), but it could also act as a distractor which would compete with drug related behavior. Interestingly, in both humans and rats, the presence of a peer effects are modulated depending on the relationship between the subject and the observer. Indeed, the presence of a stranger observing peer decreases more the cocaine consumption than the presence of a familiar one, in both rats and humans (Giorla et al., 2018). However, some questions remain to be addressed regarding the influence of socio-affective factors on behavior, such as the sensory modalities that come into play and their neurobiological basis.

Among the sensory modalities, the ultrasonic vocalizations (USV) have important communicative functions and serve as socio-affective signals in rats (Brudzynski, 2005, 2013; Nyby & Whitney, 1978). The USV of adult rats are classified into 2 major types relying on their acoustic features and emotional content: positive and negative USV (Brudzynski, 2005). Reflecting different communication functions, both types of USV elicit distinct biological and behavioral responses in recipient rats (Brudzynski, 2005, 2013; Demaestri et al., 2019).

The negative USV are characterized by a relatively stable sound frequency (from 18 to 33-kHz), very few frequency modulations and a time range from tens to thousands of milliseconds (Brudzynski, 2013). These 22-kHz long calls are known to be emitted generally in anxiogenic or aversive situations, such as predator exposure (Blanchard et al., 1991; Blanchard et al., 1990; Fendt et al., 2018), social defeat (Sales, 1972) or withdrawal from drug (Miczek & Barros, 1996; Mutschler et al., 2001; Mutschler & Miczek, 1998), and may be considered as an evolutionary equivalent of human crying (Brudzynski, 2019). These 22-kHz calls serve as signal for fear transmission (Kim et al., 2010). They elicit alertness/anxiety (Inagaki & Ushida, 2017) and defensive behaviors in the receiver, such as freezing, fight, behavioral inhibition or reduced locomotor activity (for review see Wöhr & Schwarting, 2013). Playback of 22-kHz USV induces a conditioned place aversion over a background noise playback, and when associated with cocaine self-administration, it also transiently increases cocaine intake (Montanari et al., 2020).

On the other hand, the positive 50-kHz USV calls are characterized by a broad frequency range (from 35 to 80-kHz), high variable frequency modulations and a very short duration (10-150ms) (Brudzynski, 2013). These USV are much more heterogenous than the 22-kHz USV and can be divided into several categories depending on their frequency modulation, from flat calls, containing very few frequency modulation to trills with a very high level of frequency modulation (Wöhr, 2018). Considered as an evolutionary equivalent of human laughter and joy expression (Panksepp, 2005), 50-kHz USV are emitted in appetitive situations such as tickling by a human experimenter (Panksepp & Burgdorf, 2000) or self-administration of psychostimulants (for review see Barker et al., 2015). They serve as pro-social affiliative calls (Brudzynski, 2013). Their playback is well known to induce approach behavior in recipient rat (Wöhr & Schwarting, 2007, 2009, 2012) and could also serve to orchestrate more complex social behavior, like cooperation (Łopuch & Popik, 2011). 50-kHz USV playback elicits a conditioned place preference over a background noise (Montanari et al., 2020) and when associated to cocaine self-administration, it decreases cocaine consumption (Montanari et al., 2020).

Interestingly, while there is low intra-subject variability in term of acoustic features and quantity of USV production in a same experimental context, there is a large inter-subject variability (Wright et al., 2010), depending on several factors, such as social environment, sex and strain (for review see Lenell et al., 2021). This inter-subject variability in USV suggests the existence of a vocal signature in rats, but no study to the best of our knowledge investigated the ability of rats to recognize the identity of the USV emitter.

Regarding the neurobiological basis of USV perception, it is known that replay of 50-kHz USV activates cerebral structures involved in the rewarding circuit, such as the prefrontal cortex, nucleus accumbens, thalamic parafascicular and paraventricular nuclei (Sadananda et al., 2008; Willuhn et al., 2014) and decreases the activity of the amygdala (Parsana, Li, et al., 2012). Playback of 22-kHz activates brains area associated with in fear and anxiety processing, such as perirhinal cortex, amygdala, periaqueductal grey and the antero-dorsal nucleus of the bed nucleus of the stria terminalis (Demaestri et al., 2019; Ouda et al., 2016; Parsana, Moran, et al., 2012; Sadananda et al., 2008). Less is known regarding the neurobiological substrate of the behavioral outcome of USV perception. The subthalamic nucleus (STN), a deep cerebral structure at the interface of limbic and reward circuits (Baunez & Lardeux, 2011), could play a key role. Indeed, STN inactivation differentially affects cocaine and food motivation (Baunez et al., 2005; Rouaud et al., 2010) and modulates affective responses to emotional stimuli (Pelloux et al., 2014). Moreover, STN lesion blunts the effect of 50- and 22kHz USV playback emitted by stranger rats on cocaine consumption (Montanari et al., 2020).

In this study, we first evaluated the ability of rats to recognize the USV emitter, testing if they could self-administer playback of 50- and 22-kHz USV recorded from a stranger or a familiar (i.e: their cagemate) congener. Since rats lever press to listen to 50-kHz USV emitted by strangers (Burgdorf et al., 2008) and naturally prefer novel over familiar social stimuli (Engelmann et al., 1995; Smith et al., 2015; Thor & Holloway, 1982; Veenema et al., 2012), we hypothesized that they would prefer to lever press for 50-kHz USV from the stranger rat than a familiar one if they can discriminate them. In a second experiment, we evaluated the influence of the familiarity of the USV emitter on the receiver’s behavior, testing whether the modulation induced by USV playback on cocaine consumption is a function of the familiarity of the USV emitter. We therefore expanded upon our previous experiment (Montanari et al., 2020), whereby we investigated the effect of contingent playback of 50- and 22-kHz USV on cocaine self-administration, and added the factor of familiarity of the USV emitter (stranger or home-cage partner). Finally, to investigate the neurobiological basis of this influence, we tested the effect of STN lesions on these processes.

## Material and methods

### Animals

Because female rats do not show a social novelty preference (Veenema et al., 2012) and emit less USV than males (for review see Lenell et al., 2021), this experiment was exclusively conducted on male rats. 76 male Lister Hooded rats (Charles River Laboratories, Saint-Germain-sur-l’Arbresle, France) weighing ~315 g at their arrival were used. Animals were housed in pairs, maintained in animal facility under a 12-hour inverted light/dark cycle, with all experiments conducted during the dark cycle (7am-7pm). Water and food (Scientific Animal Food and Engineering, Augy, France) were available ad libitum in the home cages. Rats were handled every day. All animal care and use conformed to the French regulation (Decree 2010-118), were approved by the local ethic committee and the French Ministry of Agriculture under the license #3129.01 and followed the 3R European rules.

### Surgery

Once rats reached ~400 g, they were anesthetized with ketamine (Imalgene, Merial, 100 mg/kg, i.p.) and medetomidine (Domitor, Janssen 30 mg/kg, i.p.) following a preventive antibiotic treatment with amoxicillin (Duphamox, LA, Pfizer, 100 mg/kg, s.c.) and placed in the stereotaxic frame (David Kopf apparatus) for bilateral injection of either 0.5 μL of 53 mM ibotenic acid (9.4 mg/mL; STN-lesioned group, n=42) or vehicle solution (phosphate buffer, 0.1 M; sham control group, n=34) into the STN (with tooth bar set at −3.3 mm; at the following coordinates anterior/posterior = −3.7 mm; lateral = ±2.4 mm from bregma; dorsoventral = −8.35 mm from skull; from Paxinos & Watson, 2007; (Paxinos & Watson, 2007) as previously described (Baunez et al., 2005; Pelloux et al., 2014). At the end of the surgical procedure, anesthesia was reversed with an injection of atipamezole (Antisedan, Janssen, 0.15 mg/kg, i.m.). For analgesia, rats were administered with meloxicam (Metacam, Boehringer Ingelheim, 1 mg/kg, s.c.) at the time of surgery.

42 rats were also subjected to intra-jugular implantation of a catheter: using standard surgical procedures (Baunez et al., 2005), silicon catheters were inserted into the right jugular vein and exited dorsally between the scapulae. The catheters were then flushed daily with a sterile saline solution containing heparin (Heparin Sodium, Sanofi, 3g/L) and enroflorilexine (Baytril, Bayer, 8g/L) during all the experiment to maintain their patency and to reduce infection risk. Catheters were also regularly tested with propofol (Propovet, Abbott, 10 mg/ml) to confirm their patency. Rats were allowed 10 days for recovery. Then they were tested for their auditory capacities, using startled auditory reflex, and their USV have been recorded before experimentation began (see below Acoustic stimuli).

### Apparatus

Standard self-administration chambers (Camden Instruments, length: 24 cm, width: 25 cm, height: 26 cm) placed in sound and light attenuating cubicles were used. Each chamber was equipped with a white house-light in the ceiling, a cue-light above each of the two retractable levers on the right-hand wall and an ultrasonic loudspeaker (Ultrasonic Dynamic Speaker Vifa, Avisoft Bioacoustics, Glienicke, Germany), connected to a portable ultrasonic power amplifier (Avisoft Bioacoustics), linked to a computer audio interface (Quad-Capture USB 2.0 Audio Capture, Roland Corporation, Los Angeles, CA, U.S) in an opening in the middle of the chamber ceiling. All the chambers described above were controlled by a custom-built interface and associated software (built and written by Y. Pelloux).

### Experimental procedures

#### 1) USV self-administration

After 5 days of habituation (i.e. animals were placed daily in the self-administration chambers for 30-min, with the levers retracted and no house-light), 34 rats were daily exposed to a 30-min session of USV playback self-administration. At the start of the session, the house-light was turned on and the two levers were extended. One press on one of the two levers delivered the playback of a USV file (1.5 sec length of either 50 or 22 kHz USV; representative files are illustrated in **Figure 1A,B)** while one press on the other lever resulted in the playback of the background-noise file (1.5 sec length; file is illustrated in **Figure 1C**). Assignation of USV or background-noise to one of the two levers were counterbalanced between animals. Each lever press switched on the cue light above the corresponding lever during 5-s, followed by a 20-s time-out period, during which the house-light was switched off and any further lever press was recorded as perseveration but had no consequence. This time-out period allowed us to discriminate between presses produced to listen to the acoustic stimulus from presses produced in reaction to the stimulus. The experiment consisted in 3 different blocks of 5 consecutive days in the following order: 50-, 22- and 50-kHz USV, .vs background-noise. The addition of the 2nd 50-kHz USV playback block allowed us to evaluate if a decrease in the number of lever presses on the USV-associated lever during the 22-kHz USV playback would be due to a general loss of interest for the task or if it could be reversed with the appetitive 50-kHz USV. Two different groups of animals were exposed to either USV emitted by a familiar (n=17, familiar USV condition) or a stranger rat (n=17, stranger USV condition). Then, a 4^th^ block of 5 consecutive days was added, during which one press on one of the two levers delivered the playback of 50-kHz USV emitted by a stranger peer, while one press on the other lever resulted in the playback of 50-kHz USV emitted by the cage-mate. Levers attribution was randomly assigned (i.e. for half of the rats, pressing on the lever formerly associated with the playback of 50- and 22-kHz USV now delivered familiar 50-kHz USV while a press on the other lever led to the playback of stranger 50-kHz USV. The opposite rule applied to the other half of the animals).

**Figure 1:**
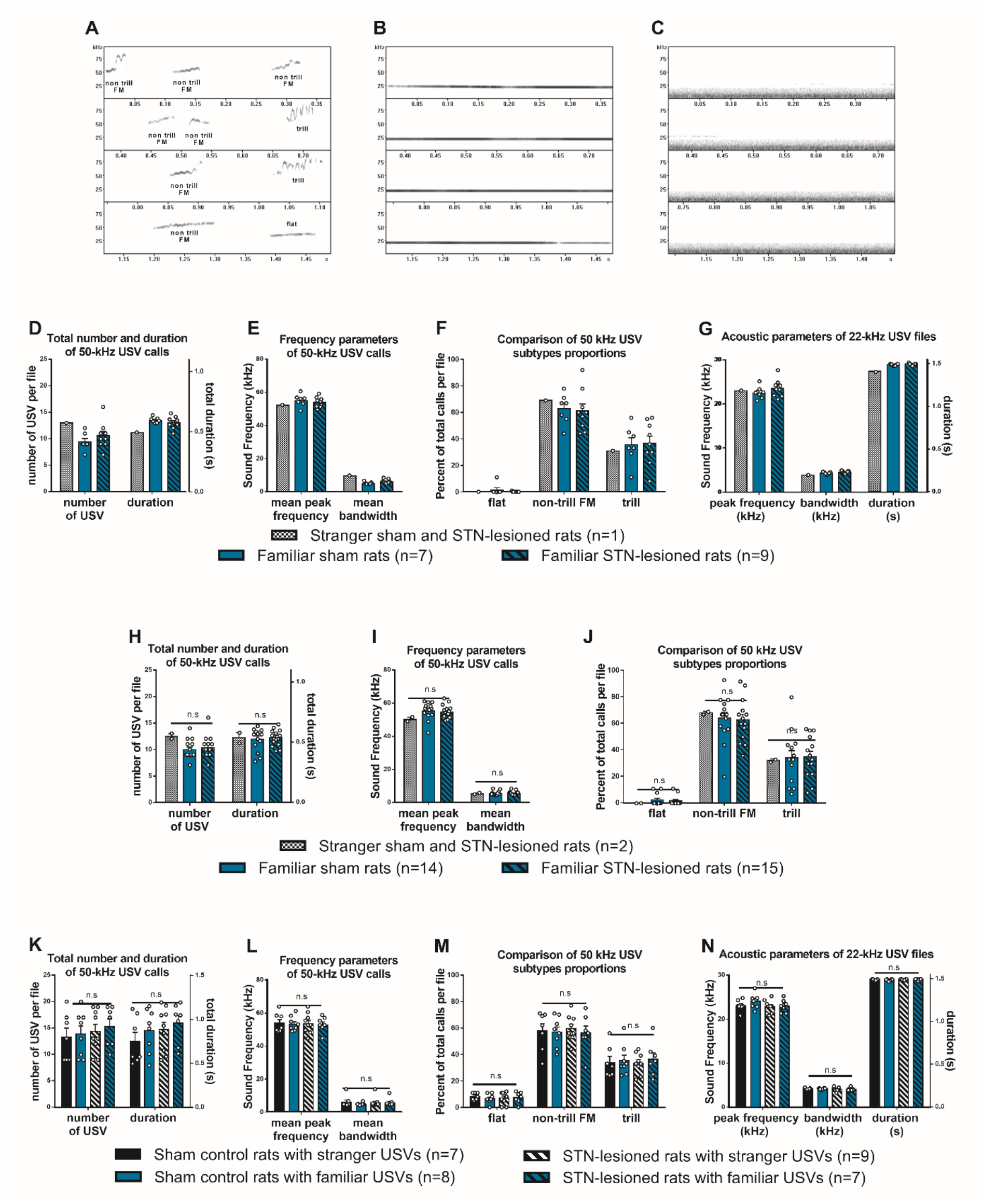
Description of the USV files. Upper graphs: Representative spectrograms with frequency (kHz) and duration (s) of the various audio stimuli used: 50-kHz USV calls (A), 22-kHz USV (B) and background noise (C). Graphs (D), (E), (F) and (G) illustrate the total number (when appropriate) and duration of the calls per file, their frequency parameters and the 50-kHz subtypes proportion of the USV files used for the 3 first blocks of the USV self-administration experiment. Graphs (H), (I) and (J) illustrate the total number and duration of the USV calls per file, their frequency parameters and subtypes proportions of the 50-kHz USV files used for the 4^th^ block of the USV self-administration experiment. Graphs (K), (L), (M) and (N) illustrate the total number (when appropriate) and duration of the calls per file, their frequency parameters and the 50-kHz subtypes proportion of the USV files used for the cocaine self-administration experiment. *FM: frequency-modulated, n.s: non-significative*

#### 2) Playback display of familiar or stranger USV during cocaine self-administration

42 other rats were trained to press a lever to self-administer cocaine for one-hour daily sessions. Before the start of the session, catheters of rats were connected to cocaine syringes positioned on motorized pumps (Razel Scientific Instruments, St-Albans, VT, USA), via infusion lines and liquid swivels. At the start of the session, the house-light was turned on and the two levers were extended. Cocaine was assigned to one of the two levers (“active lever”) and counterbalanced between rats. Pressing on the active lever delivered an intravenous infusion of cocaine (250μg per 90μl infusion in 5s) and switched on the cue-light above the active lever during the cocaine delivery. Each injection was followed by a 20-s time-out period, during which the house-light was switched off and any further lever press was recorded as perseveration but had no consequence. This time-out period was used to prevent any possible cocaine overdose. Pressing on the other lever (“inactive lever”) was recorded but had no consequence. After a few sessions ran under a continuous schedule of reinforcement (FR1; i.e. every lever press on the active lever was reinforced), the number of lever presses required to obtain a cocaine injection was increased progressively over days to FR5 (the 5th lever press delivered the infusion of cocaine). Once rats showed less than 25% of variability in the number of cocaine injections for 5 consecutive days – constituting their baseline cocaine consumption - they were exposed to the playback display during cocaine self-administration. During these playback sessions, the first lever press on the active lever resulted in the playback of the appropriate file (1.5 sec length of either 50 or 22 kHz USV or background-noise) while the 5th press started the cocaine injection. Likewise, acoustic stimulus served as preventive/incentive stimulus, associated with drug intake. These playback conditions were run in 3 different blocks of 5 consecutive days in the following order: 50-kHz USV, background-noise and 22-kHz USV. The order of the USV (50- or 22-kHz) was counterbalanced between rats. Two different groups of animals were exposed to either USV emitted by a familiar rat (familiar USV condition, i.e. cage-mate; n=21) or a stranger rat (living in a different home cage) (stranger USV condition; n=21).

### Acoustic Stimuli

USV displayed during the experiments were recorded from the tested rats at a sample rate of 192 kHz (16 bits format), using a condenser ultrasound microphone (CM16/CMPA-48, Avisoft Bioacoustics), connected to a computer audio interface (Quad-Capture USB 2.0 Audio Capture, Roland Corporation, Los Angeles, CA, U.S). The software Avisoft-SASLab Pro (Version 4.2, Avisoft Bioacustics) was used to record and mount USV files.

Long 1.5 s 22-kHz USV calls were recorded from each rat after received a mild electric foot shock (1 mA for 2 s) in a different self-administration chamber than these used for the self-administration experiments described here and located in another room to prevent negative association. For the playback of the 50-kHz USV, a series of 50-kHz calls was recorded from each rat while being tickled alone in its home cage. Background noise was extracted from one of these recordings and was always the same for all the experiments. Representative spectrograms of background noise, 50- and 22-kHz USV files used are illustrated in **Figure 1A,B,C**.

In the experiment of USV playback self-administration, for the “familiar USV condition”, every familiar USV file was collected from the cage-mate of the tested individual. The material used (50- and 22-kHz USV) for the “stranger USV condition” was recorded from a stranger rat of another experiment. Parameters of the files used for the 3 first blocks of USV playback self-administration are illustrated in **Figure 1D,E,F**, (for 50-kHz USV files used in 1^st^ and 2^nd^ 50-kHz USV playback blocks) and in **Figure 1G** (for 22-kHz USV files). For the 4^th^ block of USV playback self-administration, 2 files were played back to rats for the “stranger 50-kHz USV”. The first file, described before, was used for the rats previously self-administering familiar USV. However, to avoid an habituation effect, we played back another file to the rats previously lever pressing for stranger USV. This new file was recorded from another rat stranger to the experiment. Parameters of these 50-kHz USV files used in the 4th block are illustrated in **Figure 1H,I,J**.

For the experiment of USV playback during cocaine self-administration, each USV file served twice: first to the cage-mate of the recorded rat in the “familiar USV condition” and second, to a stranger rat in the “stranger USV condition”. Parameters of the 50- and 22-kHz USV files used in the cocaine self-administration are illustrated respectively in **Figure 1K,L,M** and **Figure 1N**.

At the end of the experiment, statistical analyses were performed, when appropriate, to detect differences between parameters (number and total duration of calls per file, mean bandwidth and peak frequency, and USV subtypes proportions) of USV files played back to the different experimental groups (see **Supplementary material**). Notably, files were similar between groups for each experiment. It is important to note that in both stranger and familiar USV playback conditions, rats were exposed to USV emitted by a sham-control or a STN-lesioned rat (in a counterbalanced way). Then, if USV playback differently affects sham-control and STN-lesioned rats, it cannot be due to a possible effect of the STN lesion on the USV emission (e.g. frequency, fluency or else), but only an effect of this STN lesion on the listening of USV playback.

As previously described (Montanari et al., 2020), we played back these acoustic stimuli at a sampling rate of 192 kHz (16 bits format). Loudspeakers had a frequency range of 1 to 120 kHz. The dB SPL was calibrated using a Calibrated Reference signal Generator (Avisoft Bioacoustics) and we used an intensity of ~70 dB SPL for the USV, corresponding to the intensity of a USV measured at 20 cm of a rat (Brudzynski et al., 1993) and ~50 dB SPL for the background noise, corresponding to its moderate intensity in the USV files.

### Histology

At the end of the experiments, rats were deeply anesthetized with isoflurane and then euthanized using pentobarbital sodium (Dolethal, Vetoquinol, 200mg/kg, intracardiac). Brains were removed and frozen in isopentane (Sigma-Aldrich) and kept at −80°C before being coronally sectioned at 40 μm thickness, using a cryostat, and stained with thionine (Sigma-Aldrich). The location and extent of STN lesions have been estimated by visualizing neuronal loss and associated gliosis. Representative intact and lesioned STN are illustrated in **Figure 2**.

**Figure 2:**
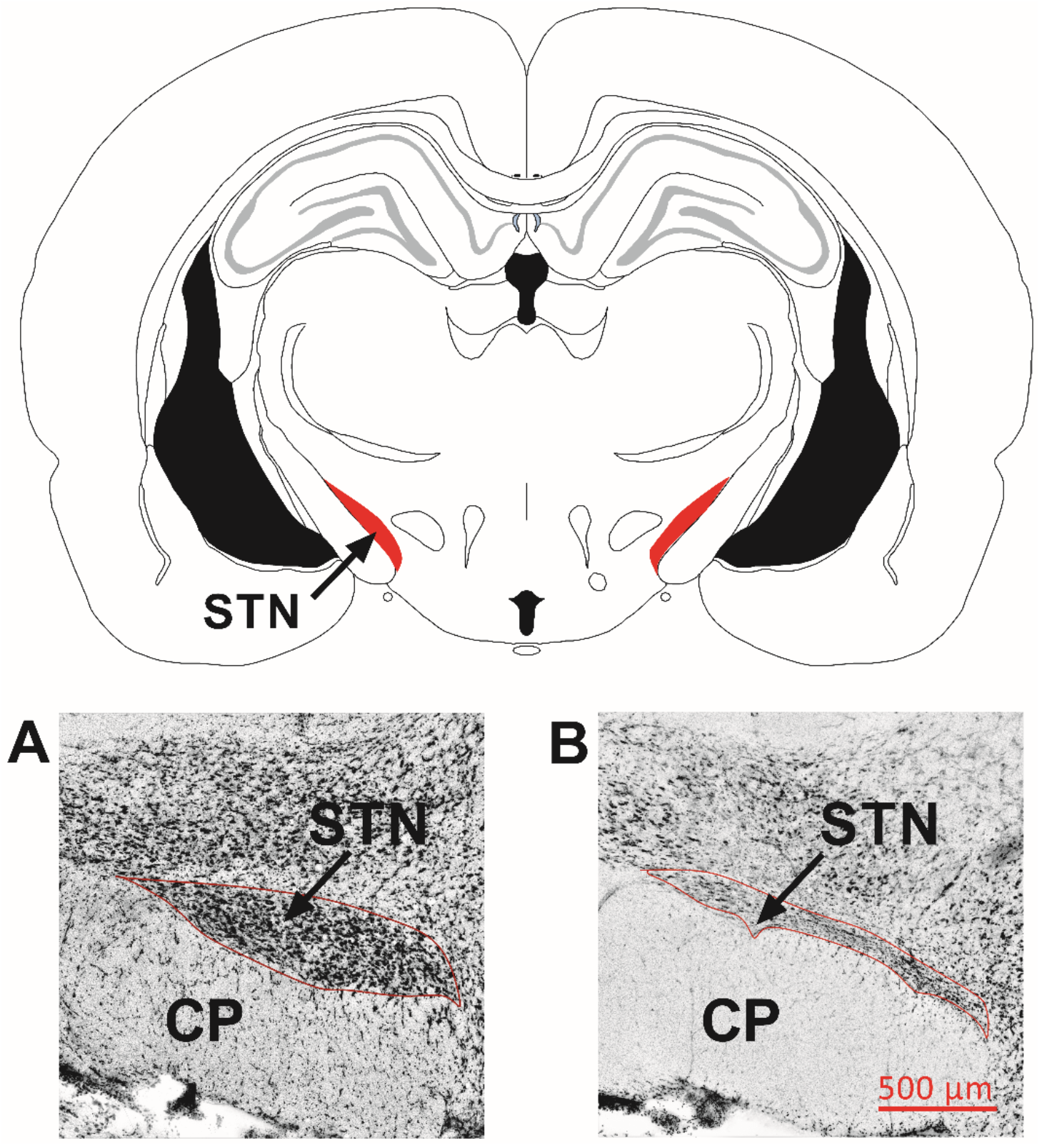
Frontal sections of the subthalamic nucleus stained with thionine. Upper panel: Schematic coronal section of the rat brain at the level of the STN targeted for the surgery (AP: −3.7 mm from bregma) (left STN in red indicated by the arrow), from Paxinos & Watson (2007). Bottom panels: Representative STN delineated by the red lines in a sham operated rat (A) and a STN-lesioned rat (B). The lesions were characterized by neuronal loss, shrinkage of the structure and gliosis. The red bar indicates the scale. *CP: cerebral peduncle; STN: subthalamic nucleus*.

### Statistical analysis

All variables are expressed as mean number ± SEM and the p-value threshold have been set at α=0.05.

#### 1) USV self-administration

Total number of presses on each lever (USV- vs background-noise-associated) was analyzed separately for sham-control and STN-lesioned rats. First, for each group (familiar and stranger USV conditions), we compared the average number of presses on each lever for the 3 playback blocks (1st 50-, 22- and 2nd 50-kHz USV, .vs background-noise) using Wilcoxon matched-pairs signed-ranks tests. We also compared the effects of these blocks on the average number of presses, separately for the USV- and background noise-associated levers, using a Friedman test. Then, we compared the average number of presses between rats self-administering familiar vs stranger USV, in each playback block, using Wilcoxon sum rank tests, separately for the two levers. To interpret these previous effects, we estimated a panel data random-effects (RE) model with the effects of USV familiarity condition, playback blocks and sessions (as dummy variables) effects on the discrimination between the two levers (i.e. number of presses on the USV-associated lever minus number of presses on the lever delivering background-noise). We adopted a panel data approach because it allows an analysis of cross-sectional time-series data. RE model controls for unobserved heterogeneity among the individuals and allows to include panel-invariant variables, such as the USV familiarity condition, among regressors. In this model, a constant contains the “reference individual”, corresponding to the discrimination of a rat subjected to stranger USV playback, on the 1st session of the 1st 50-kHz USV playback block. The parameters of the dependent variable (USV familiarity condition, playback block and session) were therefore compared to the reference individual’s parameters. In this model, a significant parameter means that the behavior diverged in a significant manner with regards to this parameter. For the analysis of the 4^th^ block of this experiment, we compared the mean number of lever presses on the familiar vs the stranger USV-associated lever, separately for the sham and the STN-lesioned rats, using Wilcoxon matched-pairs signed rank tests.

#### 2) Playback display of familiar or strangers USV during cocaine self-administration

Results were analyzed separately for sham-control and STN-lesioned rats. Because of the interindividual variability in cocaine consumption during baseline, consumption per session was divided by the average number of cocaine injections per session during the baseline, separately for each rat. First, we compared this variation ratio of cocaine consumption between each playback block (50- and 22-kHz USV, and background-noise) with the baseline via a Wilcoxon matched-pairs signed-ranks test, separately for the familiar and stranger USV conditions. Then, variation in consumption was compared between familiar and stranger USV conditions, using a Wilcoxon rank sum test, for each playback block. Finally, we estimated a panel data random-effects (RE) model with the effects of USV familiarity condition, playback blocks and sessions (as dummy variables) on the log-transformed cocaine consumption variation. Log-transformation of drug consumption with dummy variables for treatments allows us to interpret the effects as a percentage variation of consumption. In this model, the reference is the variation of cocaine consumption of a rat subjected to familiar USV playback, on the 1st session of its baseline.

The statistical tests were two tailed and were performed using R software. The graphs were performed using GraphPad Prism 6 (version 6.07).

## Results

In total, 29 rats in the analysis of USV self-administration (corresponding to n=7 stranger and n=7 familiar sham-control, and n=6 stranger and n=8 familiar STN-lesioned rats for the 3 first playback blocks, and n=14 sham-control and n=15 STN-lesioned rats for the 4th playback block) and 31 rats were included in the analysis of the effects of USV playback on cocaine consumption (corresponding to n=7 stranger and n=8 familiar sham-control, and n=9 stranger and n=7 familiar STN-lesioned rats). The other animals were excluded from STN-lesioned group because of unsatisfying lesions (partially outside the STN, too restricted or unilateral; n=11), from the sham-control group because of apparent mechanic cerebral lesion (n=1) or because they showed no startled auditory reflex or due to catheter occlusions (n=4).

### 1) Do rats lever press to listen to USV?

#### Sham-control rats self-administer stranger 50-kHz USV

As shown in **Figure 3A**, sham-control rats self-administering stranger USV playback (n=7) showed a significant preference for the lever delivering USV compared to the background-noise associated-lever, during the 1st (Wilcoxon matched-pairs signed-ranks tests: V=28, p=0.016) and 2nd (V=27, p=0.031) 50-kHz, but not during the 22-kHz, USV playback blocks (V=20, p=0.375). Friedman tests indicated an effect of the playback blocks on the mean number of presses on the USV-associated lever (χ^2^(2)=10.57, p=0.003), but not on the background-noise-associated lever (χ^2^(2)=1.143, p=0.620). Post hoc comparison revealed that rats pressed more on the USV associated-lever during the 1st (p<0.010) and 2nd (p=0.023) 50-kHz USV playback blocks than during the 22-kHz USV playback block, with no difference on the mean number of lever presses between the two 50-kHz USV blocks (p>0.999), suggesting that listening to stranger 50-kHz USV has strong and stable rewarding properties, contrary to the 22-kHz USV.

**Figure 3:**
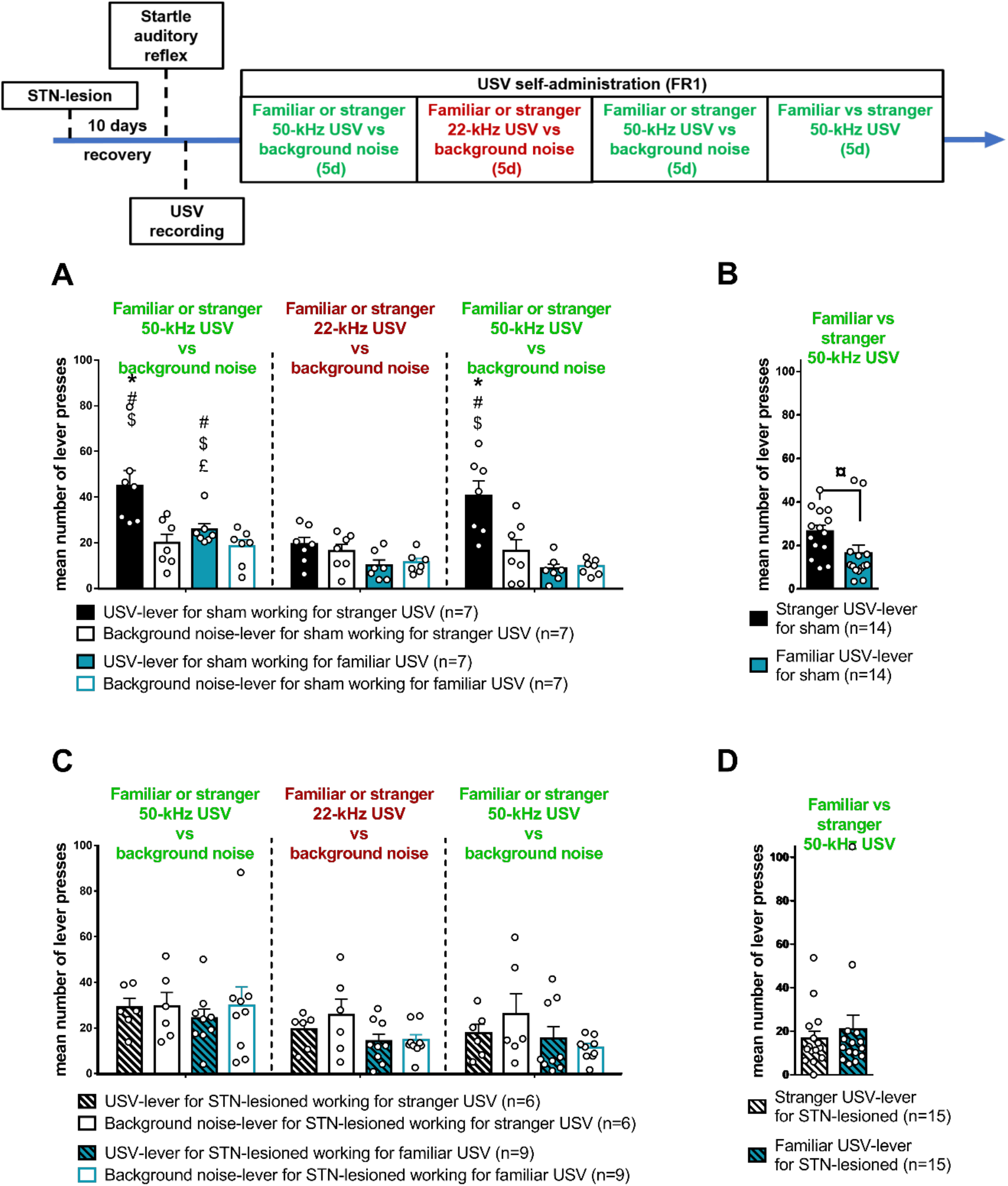
Familiar and stranger USV playback self-administration in sham-control (A and B) and STN-lesioned (C and D) rats. Upper panel: Timeline of the experiment. Bottom panels: The graphs represent the mean number of presses per session on each acoustic stimulus – associated lever during the 3 first playback blocks (1^st^: 50-kHz USV vs. background noise playback, 22-kHz USV vs. background noise playback and 2nd 50-kHz USV vs. background noise playback; A and C) and the 4^th^ playback block (familiar vs stranger 50-kHz USV; B and D) of the USV playback self-administration experiment. A: Sham-control rats subjected to stranger USV playback (n=7, black bars) pressed significantly more on USV-associated lever (full bars) during the 1st and 2nd 50-kHz USV playback blocks than on the lever delivering the background noise (empty bars) playback, but not during the 22-kHz USV playback block. Sham-control rats subjected to familiar USV playback (n=7, blue bars) pressed significantly more on the USV-associated lever (full bars) during the 1st 50-kHz USV block than on the lever delivering the background noise playback (empty bars), but not during the 2^nd^ 50-kHz USV playback block. Sham-control rats subjected to stranger USV playback pressed more on the USV-associated lever than those subjected to familiar USV playback during the two 50-kHz USV playback blocks. B: When facing the choice between a lever delivering familiar (blue bar) and stranger (black bar) 50-kHz USV, sham-control rats (n=14) showed a strong preference for the stranger USV – associated lever. C: Neither STN lesioned-rats subjected to stranger (n=6, black striped bars) nor familiar (n=9, blue stripped bars) USV playback showed a preference for one of the two levers, whatever the playback block (1st and 2nd 50-kHz, and 22-kHz USV playback). D: STN-lesioned rats (n=15) showed no preference, nor for the familiar (blue stripped bar), neither for the stranger (black stripped bar), USV – associated lever. *\*: p≤ 0.05 compared with rats submitted to familiar USV*; *#: p≤ 0.05 compared with background noise associated lever*; *$: p<0.05 compared with 22-kHz USV playback block*; *£: p<0.05 compared with the 2nd 50-kHz USV playback block*; *¤: p<0.05 compared with familiar 50-kHz USV associated lever*.

Sham-control rats self-administering familiar USV (n=7) also showed a significant preference for the lever delivering USV compared to the background-noise associated-lever, during the 1st (Wilcoxon matched-pairs signed-ranks test: V=26, p=0.031), but not the 2nd block using 50-kHz USV (V=6, p=0.688), nor during the 22-kHz USV playback block (V=17, p=0.688). Friedman test showed an effect of the playback blocks on the mean number of presses on the lever delivering USV (χ^2^(2)=10.57, p=0.003) but not on the lever delivering background-noise (χ^2^(2)=3.429, p=0.234). Post hoc comparison confirmed that rats pressed more on the lever delivering USV during the 1st 50-kHz USV playback block than during the 22- (p<0.010) and the 2nd 50-kHz (p=0.023) USV playback blocks, with no difference on the mean number of lever presses between the 22- and 2nd 50-kHz USV playback blocks (p>0.999). These results suggest that the rewarding value of the familiar 50-kHz USV observed during the 1^st^ block is blunted over time.

During the two 50-kHz USV playback blocks, rats subjected to stranger USV playback pressed more on the USV-associated lever than those subjected to familiar USV playback (Wilcoxon sum tests: 1st: W=46, p<0.004, n=14, and 2nd 50-kHz USV playback block: W=49, p<0.001, n=14), showing that stranger 50-kHz USV are more rewarding than familiar ones. Interestingly, during the 22-kHz USV block, rats lever pressing for familiar USV tended to press less on the USV-associated lever than rats self-administering stranger USV (W=40, p=0.055, n=14). In contrast, the analysis showed no difference on the number of presses on the lever delivering the background-noise between the two groups of rats, in each of the 3 playback blocks (1st: W=27, p=0.805, n=14 and 2nd 50-kHz USV: W=30, p=0.535, and 22-kHz USV: W=35, p=0.201, n=14).

As illustrated in **Figure 3B**, sham-control rats pressed significantly more on the lever delivering stranger 50-kHz USV than on the lever associated with familiar ones (Wilcoxon matched-pairs signed-ranks tests: V=85, p=0.042, n=14) confirming the preference for stranger over familiar 50-kHz USV.

Results of the RE panel data model, reported in **Table 1**, confirmed that 1) rats subjected to stranger USV playback had a stronger preference for the USV-associated lever than those subjected to familiar one during the 1st (β=−17.486, p=0.003) and the 2nd 50-kHz USV playback blocks (β=−21.857, p<0.001), 2) neither rats lever pressing for stranger nor familiar USV showed a preference for one of the two lever during the 22-kHz USV playback block (stranger: β=−21.857, p<0.001, familiar USV condition: β=−26.114, p<0.001) and 3) rats self-administering stranger USV showed a similar preference for the lever delivering USV during the 1st and the 2nd 50-kHz playback block (p=0.902). Furthermore, sessions coefficients were not significant, confirming this long-lasting effect.

**Table 1:**
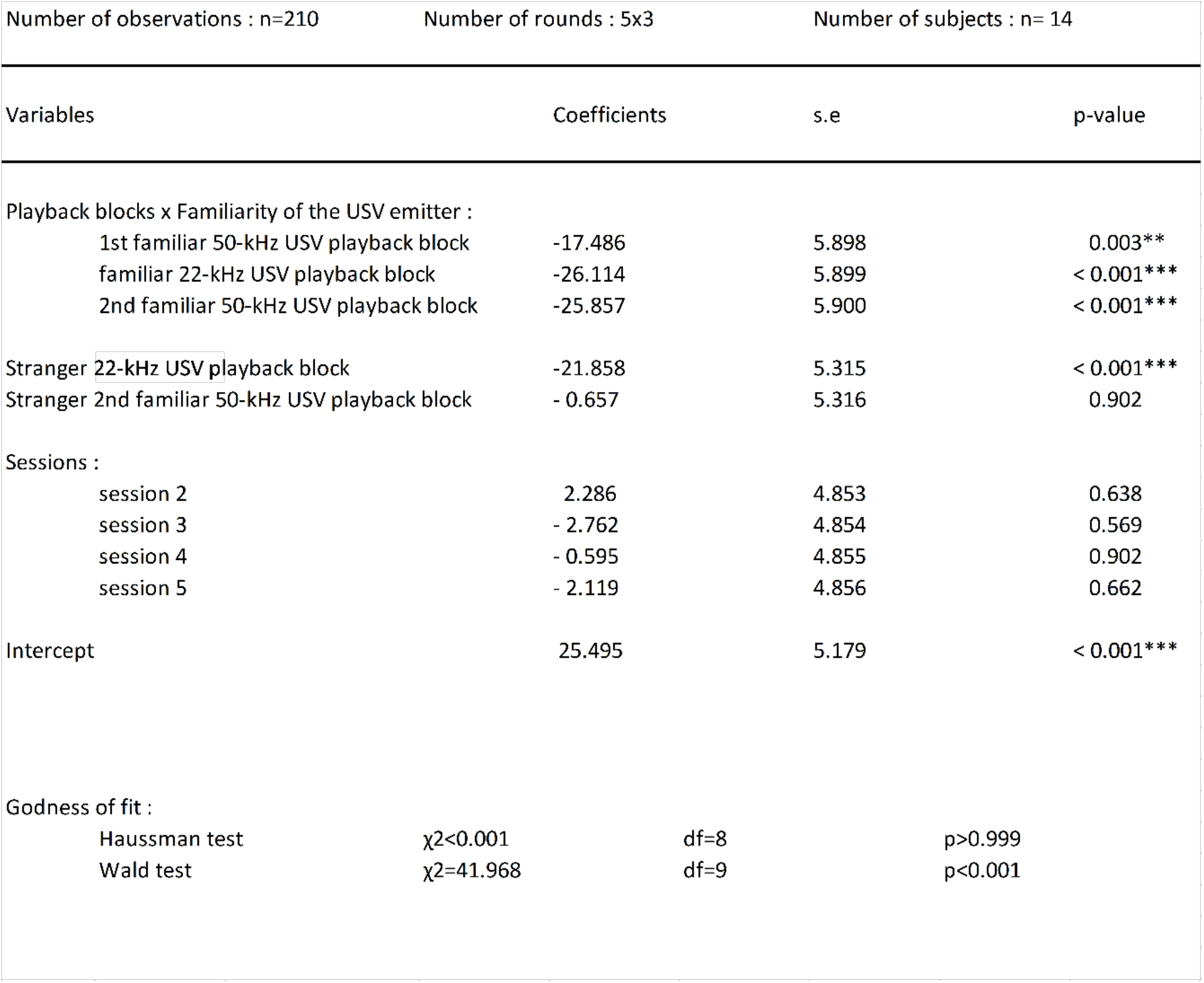
Parameters estimate for the random-effect model of sessions, playback blocks and familiarity of the USVs on the discrimination between the levers for sham-control rats. Discrimination between the two levers has been calculated as follow: number of lever presses on the lever delivering USV playback minus number of lever presses on the lever delivering background noise playback per session. In this RE model, the reference is the discrimination of a stranger rat during the 1^st^ session of the 1^st^ 50-kHz USV playback block. *\*\*: p≤ 0.005* \*\*\**: p≤ 0.001*

#### STN-lesioned rats do not press to listen to USV

As illustrated in **Figure 3C**, STN-lesioned rats self-administering stranger USV (n=6) showed no preference for one of the two levers, in any of the 3 playback blocks (Wilcoxon matched-pairs signed-ranks tests: for the 1st: V=10, p=0.590 and the 2nd 50-kHz USV: V=13, p=0.688, and for the 22-kHz: V=14, p=0.563). Besides, Friedman test only revealed a slight trend to decrease mean number of presses on the lever delivering USV across the playback blocks (χ^2^(2)= 6.333, p=0.052), and no effect on the background-noise associated-lever (χ^2^(2)=1.333, p=0.571). Then, contrary to sham-control, STN-lesioned rats subjected to stranger USV playback showed no preference for 50-kHz USV playback. These results suggest that STN-lesion blunts the rewarding effect of stranger 50-kHz USV playback.

STN-lesioned rats self-administering familiar USV playback (n=9) also showed no preference for one of the two levers, in any of the 3 playback blocks (Wilcoxon matched-pairs signed-ranks tests: for the 1st: V=17, p=0.570 and the 2nd 50-kHz USV: V=21, p=0.910, and for the 22-kHz USV: V=16.5, p=0.514). Friedman test revealed a trend to decrease the mean number of presses on the lever delivering USV (χ^2^(2)= 6, p=0.057) and on the background-noise associated-lever (χ^2^(2)=6, p=0.057) across the playback blocks. Then, contrary to sham-control, STN-lesioned rats lever pressing for familiar USV showed no preference for 50-kHz USV playback, even during the 1^st^ playback block.

Familiarity with the USV emitter had no effect on the mean number of presses on the lever delivering USV in any of the playback blocks (Wilcoxon sum tests: 1st: W=35, p=0.388, n=15 and 2nd 50-kHz USV: W=34, p=0.456, n=15, and 22-kHz USV: W=34, p=0.456, n=15), nor on the background-noise associated-lever (1st: W=39, p=0.157, n=15 and 2nd 50-kHz USV: W=38.5, p=0.194, n=15, and 22-kHz USV: W=39.5, p=0.157, n=15). These results confirmed that, contrary to sham-control, STN-lesioned rats show no sensitivity to the familiarity with the rat emitting USV. Confirming this effect, when STN-lesioned rats faced the choice between 50-kHz familiar and stranger USV playback, they showed no preference for one of the two levers (V=53, p=0.720, n=15), as shown in **Figure 3D**.

Taken together, these results on USV self-administration reveal that stranger, but not familiar, 50-kHz USV have strong and stable rewarding properties for sham-control rats and that sham-control rats strongly prefer the stranger over the familiar 50-kHz USV. Besides, the STN-lesion makes rats insensitive to USV, whatever their emotional valency and their emitter.

### 2) How the familiarity with the USV emitter modulates the effect of the USV playback on cocaine consumption?

#### Stranger, but not familiar, 50-kHz USV playback decreases cocaine intake in sham-control rats

As illustrated in **Figure 4A**, sham-control rats subjected to stranger USV playback (n=7) showed a significant reduction in their cocaine consumption between the baseline (mean number of injections: 21.03, ±3.33) and the 50-kHz USV playback block (Wilcoxon matched-pairs signed-ranks test: V=28, p=0.016). In contrast, no difference was found under background-noise condition when compared with the baseline (V=17, p=0.688), nor when rats were exposed to 22-kHz USV playback (V=21, p=0.297). Nevertheless, a trend modulation towards increased cocaine consumption between the 1st and 5th session of the 22-kHz USV playback was observed (Wilcoxon matched-pairs signed-ranks: V=1, p=0.0592). The fact that the emotional valence of the USV emitted by a stranger rat, played back during cocaine self-administration, differently affects the drug consumption is in line with former results (Montanari et al., 2020).

**Figure 4:**
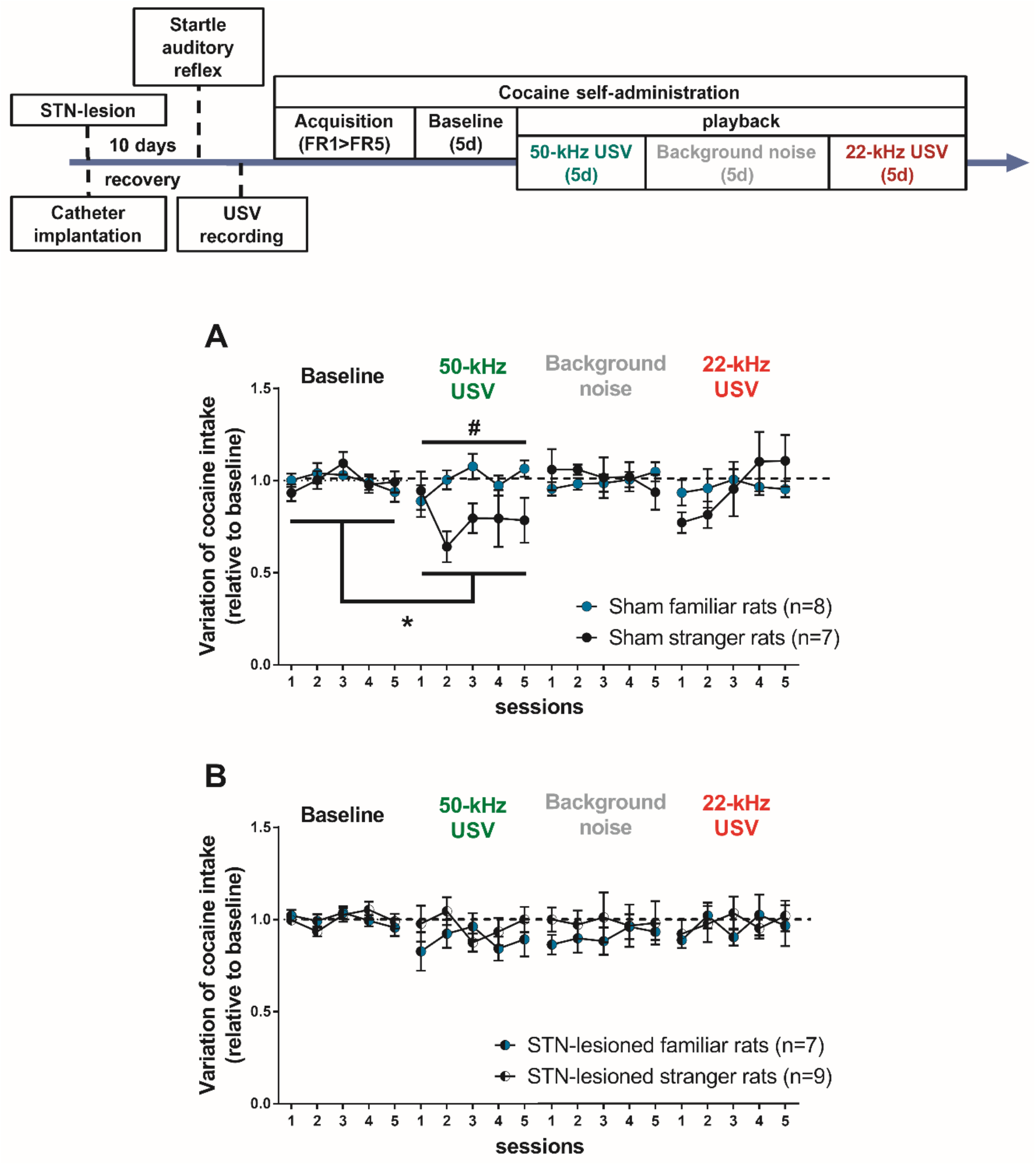
Variation of cocaine consumption under familiar and stranger 50- and 22-kHz USV and background noise playback in sham-control (A) and STN-lesioned (B) rats. Upper panel: timeline of the experiment. Bottom panels: The graphs represent the mean variation of cocaine intake relative to baseline (± SEM) for sham (A; n=15; submitted to stranger USVs condition, n=7, black dots; to familiar USVs condition, n=8, blue dots) and STN-lesioned rats (B; n=16; submitted to stranger USVs condition, n=9, black and white dots; to familiar USVs condition, n=7, blue and black dots) during the 5 sessions of each playback block (50- and 22-kHz USVs and background noise). The dotted line represents the 1 value in variation, i.e. no change from baseline. *\*: p≤ 0.05 compared with baseline; #: p≤ 0.05 compared with familiar USVs condition*.

In contrast, no difference in cocaine consumption of rats subjected to familiar USV playback (n=8) was been found between the baseline (mean number of injections: 15.18, ±2.02) and any playback blocks (Wilcoxon matched-pairs signed-ranks tests: baseline vs 50-kHz USV: V=20, p=0.844, vs background-noise: V=22, p=0.641 and vs 22-kHz USV: V=25, p=0.383), suggesting that emotional valence of USV emitted by a familiar rat does not modulate the consumption. Furthermore, cocaine intake significantly differed between rats subjected to stranger vs familiar USV during 50-kHz USV playback (Wilcoxon rank sum test: W=9, p=0.0289, n=15), but not during background-noise (W=31, p=0.779, n=15), nor 22-khz USV (W=21, p=0.463, n=15) playbacks.

As a robustness check, results from the RE panel data model, reported in **Table 2**, confirmed that stranger (β=−25%, p=0.003), but not familiar (p=0.452), 50-kHz USV playback strongly decreased cocaine consumption. Familiarity status with the USV emitter had no general effect (p=0.922), thereby specifically regulating the effect of 50-kHz USV on cocaine consumption. Furthermore, no effect of the sessions was found, suggesting that there is no habituation within the playback blocks.

**Table 2:** Parameters estimate for the random-effect model of sessions, playback blocks and familiarity of the USVs on the log-transformed cocaine consumption of sham-control rats. In this RE model, the reference is the consumption of a rat subjected to familiar USV playback during the 1st session of the baseline. *\*\*: p≤ 0.005*

| Number of observations : n=300 |  | Number of rounds : 5x4 |  | Number of subjects : n= 15 |  |
| --- | --- | --- | --- | --- | --- |
| Variables |  | Coefficients |  | s.e | p-value |
| Playback blocks : |  |  |  |  |  |
| 50-kHz USV |  | -0.044 |  | 0.059 | 0.0645 |
| background noise |  | 0.016 |  | 0.058 | 0.787 |
| 22-kHz USV |  | -0.095 |  | 0.058 | 0.102 |
| Familiarity of the USV emitter |  | -0.007 |  | 0.075 | 0.922 |
| Stranger 50-kHz USV playback |  | -0.250 |  | 0.085 | 0.003** |
| Stranger background noise playback |  | -0.034 |  | 0.085 | 0.684 |
| Stranger 22-kHz USV playback |  | 0.050 |  | 0.085 | 0.557 |
| Sessions : |  |  |  |  |  |
| session 2 |  | -0.005 |  | 0.047 | 0.912 |
| session 3 |  | 0.061 |  | 0.047 | 0.195 |
| session 4 |  | 0.300 |  | 0.047 | 0.530 |
| session 5 |  | -0.026 |  | 0.047 | 0.576 |
| Intercept |  | -0.027 |  | 0.059 | 0.645 |
| Godness of fit : |  |  |  |  |  |
| Haussman test |  | χ2=1.280 | df=10 | p=0.257 |  |
| Wald test |  | χ2=38.071 | df=11 | p<0.001 |  |

#### Cocaine consumption of STN-lesioned rats is not affected by USV playback

As illustrated in **Figure 4B**, no variation of cocaine consumption was observed compared with the baseline (mean number of injections: 17.87, ±3.35) for the STN-lesioned rats subjected to stranger USV (n=9) (Wilcoxon matched-pairs signed-ranks tests: baseline vs 50-kHz USV: V=20, p=0.353, vs background-noise: V=22, p>0.999, vs 22-kHz USV playback: V=27, p=0.652), in line with former results (Montanari et al., 2020). STN-lesioned rats subjected to familiar USV (=7) also showed no variation in their consumption compared with the baseline (mean number of injections: 16.63, ±4.62) (Wilcoxon matched-pairs signed-ranks tests: baseline vs 50-kHz USV: V=22, p=0.219, vs background-noise: V=23, p=0.156, vs 22-kHz USV playback block: V=18, p=0.578). Furthermore, the familiarity with the USV’s emitter showed no modulation of cocaine consumption, in any playback block (Wilcoxon rank sum tests: 50-kHz USV: W=16, p=0.112, n=15, background-noise: W=24, p=0.470, n=15, and 22-kHz USV: W=30, p=0.918, n=15).

## Discussion

Here, we have shown that in sham-control rats, 50-kHz USV playback was more rewarding if emitted by a stranger than a familiar rat. Consequently, only stranger 50-kHz USV playback decreased cocaine intake. Independently of the USV emitter, rats did not work to listen to 22-kHz USV and their playback had no effect on cocaine consumption. In contrast, neither the familiarity nor the emotional valence of the USV playback affected the STN lesioned rats which, therefore, did not lever press to listen to their playback and remained stable in their cocaine intake whatever the playback condition.

### Effect of the familiarity on the rewarding properties of USV and consequences on cocaine intake

Here, we have shown that rats actively lever press for stranger 50-kHz USV playback, in line with former work (Burgdorf et al., 2008), confirming the rewarding properties of those USV (Burgdorf et al., 2008; Montanari et al., 2020). Moreover, the rewarding properties of stranger 50-kHz USV playback remained stable over the sessions, since rats continuously pressed more on the lever associated with USV than on the background-noise associated lever. On the other hand, rats subjected to familiar USV playback pressed less on the USV-associated lever than animals subjected to stranger one during both 50-kHz USV playback blocks. Furthermore, their preference for these USV over the background noise extinguished over time and is no longer significant during the 2^nd^ 50-kHz USV playback block. This changed preference could rely on the fact that rats are probably habituated to 50-kHz calls from their cage-mates. Finally, when facing the choice between stranger and familiar 50-kHz USV, rats lever pressed more to listen to stranger than familiar USV. Taking together, results showed that listen to stranger 50-kHz USV is more rewarding than listen to familiar ones, which is in line with rats’ social novelty preference (Engelmann et al., 1995; Smith et al., 2015; Thor & Holloway, 1982; Veenema et al., 2012).

This preference of the sham-control rats for stranger over familiar 50-kHz USV cannot be explained by differences between groups in USV files parameters (subtypes composition, total duration and number of calls, mean peak frequency and bandwidth), since they are similar between rats subjected to stranger and familiar USV. However, in accordance with literature (Lenell et al., 2021; Wright et al., 2010), we observed an inter-subject variability in these USV files parameters within groups. Then, this differential preference for familiar vs stranger USV indicates that rats are able to recognize the vocal signature of their cage-mate.

The fact that stranger 50-kHz USV playback diminished cocaine consumption in our experiment is in line with previous results (Montanari et al., 2020) and could rely on the hypothesis that social reward – here 50-kHz USV playback - can counterbalance the drug reinforcing properties (Fritz et al., 2011). Indeed rats consume less drug in the presence of a peer than when alone, and even less if this peer is a stranger (Giorla et al., 2018). Also, when they can choose between drug and social interaction, social reward induces abstinence, confirming the beneficial effect of proximal social context on drug intake (Venniro et al., 2018). Besides, both cocaine intake and listening to 50-kHz USV activates the reward system, highlighting their competing reinforcing effects. However, the influence of the familiarity of USV on this activation of the reward system has not yet been assessed.

In contrast to stranger, familiar 50-kHz USV playback did not affect cocaine consumption. It is thus highly possible that the rewarding value of familiar 50-kHz USV playback is not strong enough to counterbalance the reinforcing effect of cocaine. The lower impact of familiar USV playback (compared with unfamiliar) during cocaine self-administration is also in line with the influence of familiarity of an observing peer during drug consumption (Giorla et al., 2018). Indeed, the presence of a stranger peer decreased more the cocaine intake than the presence of the cage-mate. One could hypothesize that the poor rewarding properties of familiar presence and USV playback and their consequences on drug consumption cannot be transposed on human because rats do not choose their cage-mate, contrary to human beings, assumed to have chosen their close partners. This difference could result in a serious change in the rewarding value of familiar stimuli. However, in Giorla et al. (2018) experiment, like in rats, human stimulant users reported to take less drug in the presence of a stranger over a familiar peer. Taken all together, our results suggest that positive communication and familiarity constitute key factors in the influence of proximal social factors on cocaine consumption.

Results from the USV playback self-administration did not show an aversive effect of the 22-kHz USV when emitted by a stranger rat, since animals did not self-administer less these USV than background noise. On the contrary, we observed a trend to avoid 22-kHz USV when emitted by a familiar rat. Otherwise, neither stranger nor familiar 22-kHz USV playback did influence rats’ cocaine consumption. In contrast, in Montanari et al. (2020) experiment, stranger 22-kHz USV playback induced a conditioned place aversion procedure and a transient increase in cocaine intake, confirming that 22-kHz USV playback has aversive properties.

Discrepancies between observed behavioral responses to 22-kHz USV across studies are common (for more details see Wöhr & Schwarting, 2013) and could rely on the way different factors can interfere on the behavioral response to their playback. For instance, Burman at al. (2007) measured behavioral fear response of rats subjected to either background-noise or 22-kHz USV playbacks emitted by 2 different rats. Their results showed that 22-kHz USV emitted only by one animal, but not the other one, elicited a behavioral response significantly different to that induced by background-noise, whereas both rats were from the same strain and had approximately the same age. These results could suggest either that 1) the two different USV files convey some variations in their content (e.g. the intensity of the aversive event), or 2) the USV’s emitter identity could be an important factor for the induced behavioral reaction (e.g. hierarchical status).

Moreover, rats’ environment and anxiety history are known to modulate their fear responses (Harding et al., 2004; Yee et al., 2012). For instance, Kim et al., (Kim et al., 2010) have shown that only rats with a first exposure to stress before experiencing 22-kHz USV, but not naïve animals, expressed a fear related response to this playback. Further studies are needed to better understand this variability in behavioral responses to the 22-kHz USV calls.

It is important to note that this study was exclusively conducted on male rats, as a first step before testing female rats, although they do not seem to show social novelty preference (Veenema et al., 2012) and emit less USV than males (for review see Lenell et al., 2021). There is, to our knowledge, no study investigating the female rat behavioral response to USV emitted by female. We will try to fill this gap in the future to provide a better knowledge of rats USV and their influence on both genders.

### Effect of STN lesions on the familiarity effect, the rewarding properties of USV and consequences on cocaine intake

Our results from the USV playback self-administration indicated that STN-lesioned animals never discriminated between USV- and background-noise associated levers, whatever the emotional valency of the USV and the familiarity of the emitter. The lack of rewarding properties of the 50-kHz USV emitted by a stranger or a familiar rat in these STN-lesioned animals, is in line with former results (Montanari et al., 2020). Because STN-lesioned rats did never show a preference for one lever, whatever the emotional valence of the USV playback, we cannot conclude regarding the effect of the STN lesion on the aversive properties of 22-kHz USV emitted by a stranger or a familiar rat. However, Montanari et al. (2020) have shown that STN lesions blunt the aversive properties of stranger 22-kHz USV, in a conditioned place aversion. Furthermore, USV playback did not affect cocaine consumption of the STN-lesioned rats, whatever their emotional value or the familiarity with their emitter.

Interestingly, in human, the involvement of STN in social emotions has been documented thanks to the application of STN deep brain stimulation (DBS) in Parkinson’s disease (PD) patients. Indeed, STN-DBS induces alterations of social emotion decoding (Coundouris, 2019), notably vocal emotion (Brück et al., 2011; Péron et al., 2010). Besides, in healthy subjects, the STN has a functional connectivity with structures involved in emotional prosody decoding (Péron et al., 2016). In agreement with these human studies, our results could suggest that in rats STN lesions affect USV emotional content decoding, thereby blunting the emotion contagion induced by the listening to the USV, and in fine, the behavioral response to those USV.

STN inactivation also diminishes general affective state. Indeed, in human, STN-DBS of PD patients can induce apathy (Rodriguez-Oroz et al., 2012; Zoon et al., 2021). Likewise, in rats, STN lesion or chronic stimulation leads to deficits in emotional response to hedonic and aversive stimuli (Pelloux et al., 2014; Vachez et al., 2020). This diminished affective state could then be responsible for the blunted behavioral response to the USV. Alternatively, unpublished data of our team showed that STN lesion abolishes social recognition in rats. We cannot exclude the possibility than in STN-lesioned rats, an inability to differentiate USV emitted by a stranger rat interferes with the effect conveyed by the USV playback, thereby blunting it.

Importantly, in PD patients, STN-DBS also affects vocal expression. Indeed, it has been shown that STN-DBS could improve the oral motor control of PD patients’ speech (Gentil et al., 1999). Although in parallel, it could negatively impact their emotional prosody expression, in correlation with its decoding (Jin et al., 2017; Kastamoniti et al., 2017). These effects would depend on the exact location of the electrode’s implantation in the STN (e.g. limbic vs sensorimotor part) (Jorge et al., 2020). Besides, previous work has shown that STN-lesioned rats emit less 22-kHz USV in response to an aversive stimulus (Pelloux et al., 2014). It would be interesting for future investigations to further investigate this implication of the STN on emotional prosody expression, to better understand the differential effects of its stimulation in PD patients.

In every case, our results confirm the involvement of the STN in vocal emotion decoding and more broadly, its role on the influence of proximal social factors on drug consumption.

## Conclusions

In conclusion, rats can differentiate the vocal signature of 50-kHz USV emitted by their cage-mate, over a stranger rat. The playback of stranger 50-kHz USV has stronger and more long-lasting properties than playback of familiar ones. Those rewarding properties might counterbalance cocaine effects, leading to a reduced drug consumption when stranger 50-kHz USV playback is paired with cocaine self-administration. The STN modulates the rewarding properties of the stranger 50-kHz USV playback, and their effect on cocaine consumption. Taken as a whole, our study advocates for the existence of an identifiable vocal signature in rats, shows the importance of ultrasonic communication in socio-affective influence of behavior, notably in the influence of social proximal factors on drug consumption and confirm the role of the subthalamic nucleus on this influence.

## Acknowledgements

The authors thank Stephane Luchini for helping with the data analysis, Julien Poitreau for helping with histological work, Linda Piermatti and Pelin Ayyildiz for helping with behavioral testing, Joel Baurberg for technical assistance with all electronic devices and the technical staff of the animal facility for ensuring the well-being of our animals.

## Author Contributions

CV: designed the research, performed research, analyzed data and wrote the paper;

CM: performed the research and contributed to experimental material;

YP: designed the research and contributed to analytical tools;

CB: designed and supervised the research, performed surgery, wrote the paper and obtained the funding.

## Supplementary material: statistical analyses of the USV files

Parameters of the files used for the 3 first blocks of USV playback self-administration are illustrated in **Figure 1D,E,F**, (for 50-kHz USV files used in 1^st^ and 2^nd^ 50-kHz USV playback blocks) and in **Figure 1G** (for 22-kHz USV files). No statistical analysis was performed on these files since only one was used for all the rats exposed to the stranger USV playback.

For the 4^th^ block of USV playback self-administration (illustrated in **Figure 1G,H,I**), a Kruskal-Wallis test was performed to detect differences between the different parameters of files played back to the 3 experimental groups (i.e: files for familiar sham rats; n=14, files for familiar STN-lesioned rats; n=15 and files for stranger sham and STN-lesioned rats; n=2). The analysis revealed no difference in files between groups, regarding the total number (χ^2^=4.045, p=0.132) and duration (χ^2^=0.013, p=0.993) of calls per file, nor on their mean peak frequency (χ^2^=3.545, p=0.170) and bandwidth (χ^2^=0.115, p=0.944). Regarding the USV calls subtypes, the Kruskal-Wallis test revealed no difference neither (for flats: χ^2^=0.524, p=0.770, non-trill FM: χ^2^=0.783, p=0.676 and trill: χ^2^=0.069, p=0.966 calls proportions).

Parameters of the 50-kHz USV files used in the cocaine self-administration in **Figure 1K,L,M**. A Kruskal-Wallis test was performed to detect differences between the different parameters of files in the 4 experimental groups (i.e: files for familiar sham; n=8, files for stranger sham; n=7, files for familiar STN-lesioned rats; n=7 and files for stranger STN-lesioned rats; n=9). The analysis revealed no difference in files between groups, regarding the total number (χ^2^=1.221, p=0.748) and duration (χ^2^=2.922, p=0.404) of calls per file, nor on their mean peak frequency (χ^2^=0.055, p=0.997) and bandwidth (χ^2^=0.424, p=0.935). The Kruskal-Wallis test also showed no difference between groups in files regarding calls subtypes composition (for flats: χ^2^=0.613, p=0.893, non-trill FM: χ^2^=0.284, p=0.963 and trill: χ^2^=0.340, p=0.952 calls proportions). Finally, parameters of the 22-kHz USV files used in the cocaine self-administration are illustrated in **Figure 1N**. Again, a Kruskal-Wallis revealed no difference in files between groups regarding the total duration of the call (χ^2^=2.540, p=0.468), nor its mean peak frequency (χ^2^=2.577, p=0.462) and bandwidth (χ^2^=2.479, p=0.479).

